# The role of consciously timed movements in shaping and improving auditory timing

**DOI:** 10.1101/2022.10.24.513592

**Authors:** Rose De Kock, Weiwei Zhou, Poorvi Datta, Wilsaan Mychal Joiner, Martin Wiener

## Abstract

Our subjective sense of time is intertwined with a plethora of perceptual, cognitive, and motor functions, and likewise, the brain is equipped to expertly filter, weight, and combine these signals for seamless interactions with a dynamic world. Until relatively recently, the literature on time perception has excluded the influence of motor activity, yet, it has been found that motor circuits in the brain are at the core of most timing functions. Several studies have now identified that concurrent movements exert robust effects on perceptual timing estimates, but critically, have not assessed how humans consciously judge the duration of their own movements. This creates a gap in our understanding of the mechanisms driving movement-related effects on sensory timing. We sought to address this gap by administering a sensorimotor timing task in which we explicitly compared the timing of isolated auditory tones and arm movements, or both simultaneously. We contextualized our findings within a Bayesian cue combination framework, in which separate sources of temporal information are weighted by their reliability and integrated into a unitary time estimate that is more precise than either unisensory estimate. Our results revealed differences in accuracy between auditory, movement, and combined trials, and crucially, that combined trials were the most accurately timed. Under the Bayesian framework, we found that participants’ combined estimates were more precise than isolated estimates in a way that trended towards optimality, while being overall less optimal than the model’s prediction. These findings elucidate previously unknown qualities of conscious motor timing, and proposes computational mechanisms that can describe how movements combine with perceptual signals to create unified, multimodal experiences of time.

## Introduction

Motor control functions are critical to our survival in the world and diverse in nature, spanning multiple time scales and integrating a flood of neural signals to guide us through various tasks ((Cisek & Kalaska, 2010); (Alhussein & Smith, 2021); (Kaplan et al., 2020)). Important to movement is the monitoring of sensory information to update movement plans according to errors or environmental demands (Franklin & Wolpert, 2011). Additionally, individuals often calibrate ongoing movements to amplify or suppress channels of sensory information according to their goals via “active sensing” (Schroeder et al., 2010), reflecting the continuous and bidirectional nature of the relationship.

It follows that time perception, a high-level, cumulative evaluation of one or more sensory channels, is highly malleable in response to movement characteristics. Although time perception studies have largely excluded motor components, recent studies focused on arm movements reveal that the accuracy and precision of perceptual timing are affected by a number of movement characteristics such as direction, speed, distance, and movement environment ((Tomassini & Morrone, 2016); (Yokosaka et al., 2015); (Yon et al., 2017); (De Kock et al., 2021)a). Notably, movement can also improve time perception. In a set of complementary experiments, auditory intervals were presented either during arm movements or in the absence of movement. For both temporal categorization and reproduction tasks, intervals encoded during movement were timed more precisely (Wiener et al., 2019). In a separate temporal discrimination study using auditory intervals, timing precision was enhanced for intervals for which the stimulus onset was determined by the participant rather than passively presented (Iordanescu et al., 2013). The benefit of movement is also highlighted in animal behavior. Rats trained to estimate a fixed interval to receive a reward learned to use stereotyped movements to enhance the accuracy and precision of their estimates (Safaie et al., 2020).

Given the clear temporal benefit of movement, it is reasonable to ask whether and to what extent it is timed with different levels of accuracy and precision than the sensory channels that much of this work focuses on, and whether these differences can explain how the channels of information are combined. To address these questions, we contextualize our study under a framework of Bayesian cue combination, which posits that multiple channels of timing information are evaluated by their reliability and optimally integrated into a more precise estimate than either alone (De Kock et al., 2021). Accordingly, the mean of the combined estimates is predicted to gravitate towards the mean of the more precise (i.e., more influential) modality. This framework aligns with principles of multisensory cue combination ((Seilheimer et al., 2014); (Ma et al., 2006; Alais & Burr, 2019)), which place importance on signal reliability when combining multiple inputs. Neural data reflect this differential weighting via population responses in multisensory areas ((Fetsch et al., 2011); (Gu et al., 2008)).

This evidence has offered insights upon which to build our understanding of the intersection of motor and timing processes (Merchant & Yarrow, 2016). However, little is known about how self-movements are timed without an added perceptual event. We note that motor control studies certainly include timing components, but the key caveat is that they examine implicit rather than explicit timing. For example, a task might require participants to synchronize their movements to a beat (Repp & Su, 2013) or interact dynamically with a stimulus without probing their conscious evaluation of the passage of time. One preliminary study of interest (Guo et al., 2019) employed a unique paradigm to test explicit timing of durations that were implicitly encoded. Participants were trained on a “skittles” task requiring them to hold and release a virtual ball to hit a target. Repeated practice led them to internalize an optimal duration range for which holding and releasing led to a successful trial. When tested in an explicit timing task, they exhibited a selective improvement for timing the target interval, while participants who did not play the game showed no benefit. Combined with the previous studies discussed, this supports the hypothesis that movements offer highly reliable temporal measurements that in turn improve timing of concurrent events and even future timing performance. This is further evidenced by increased timing acuity in motor “experts” such as athletes and drummers ((Chen & Cesari, 2015); (Cicchini et al., 2012)).

Our goal in the current study was twofold: first, we sought to understand how duration of self-movement is evaluated, given that most movement-timing tasks have either focused solely on implicit timing or have not isolated movements from a concurrent perceptual event. Second, we sought to describe differences in motor and auditory timing, and importantly, how these sources of information are combined to form a unitary estimate. We found evidence that there are differences in timing accuracy between motor and auditory estimates, and that durations are both more accurately and precisely timed using both sources of information. Finally, we synthesized these results under a Bayesian cue combination framework to account for the pronounced benefit that resulted from combining motor and perceptual sources.

## Methods

### Participants

We tested twenty right-handed participants (13 female, 7 male, M age = 25.45(9.17)). Handedness was confirmed by the Edinburgh Handedness Inventory (Oldfield, 1971). Procedures were approved by the University of California, Davis Institutional Review Board.

### Procedure

Participants performed the experiment using a robotic arm manipulandum (KINARM End-Point Lab, BKIN Technologies; (Nguyen et al., 2019); (Hosseini et al., 2017)) that allowed movement along a flat workspace using the right arm. Direct viewing of the robotic arm was occluded by a flat display that allowed viewing of targets and cues via a downward-facing monitor mounted above the workspace. Motor output was sampled at 1000 Hz. Participants were free to adjust the chair so they could comfortably view the full display.

Trials were divided into encoding and reproduction phases, and were structured as follows (see Fig. 1): first, the robotic arm guided participants to one of 16 locations in a grid-like array. Then, they experienced one of three trial conditions. In “movement” trials, subjects began moving until interrupted by an imposed brake (a 100 ms linear increase in resistive force from 0 to 50 N). In “auditory” trials, the robotic arm was locked in the random location and the participant heard an auditory tone. In “combined” trials, subjects were cued to move while timing a concurrent auditory tone; in this condition, the tone began as soon as the apparatus detected movement at the velocity threshold of 5 cm/s, and the brake was applied synchronously with the auditory tone offset. After the encoding phase, they were guided to a central target for the reproduction phase. When this target turned green, subjects reproduced the encoded duration by holding and releasing a button attached to the handle. The tested durations were 1000, 1500, 2000, 2500, 3000, 3500, and 4000 ms. Trial conditions were experienced in blocks of 14 trials (for a total of 210 trials) in a pseudorandomized order such that no condition was experienced twice in a row.

**Figure 1:**
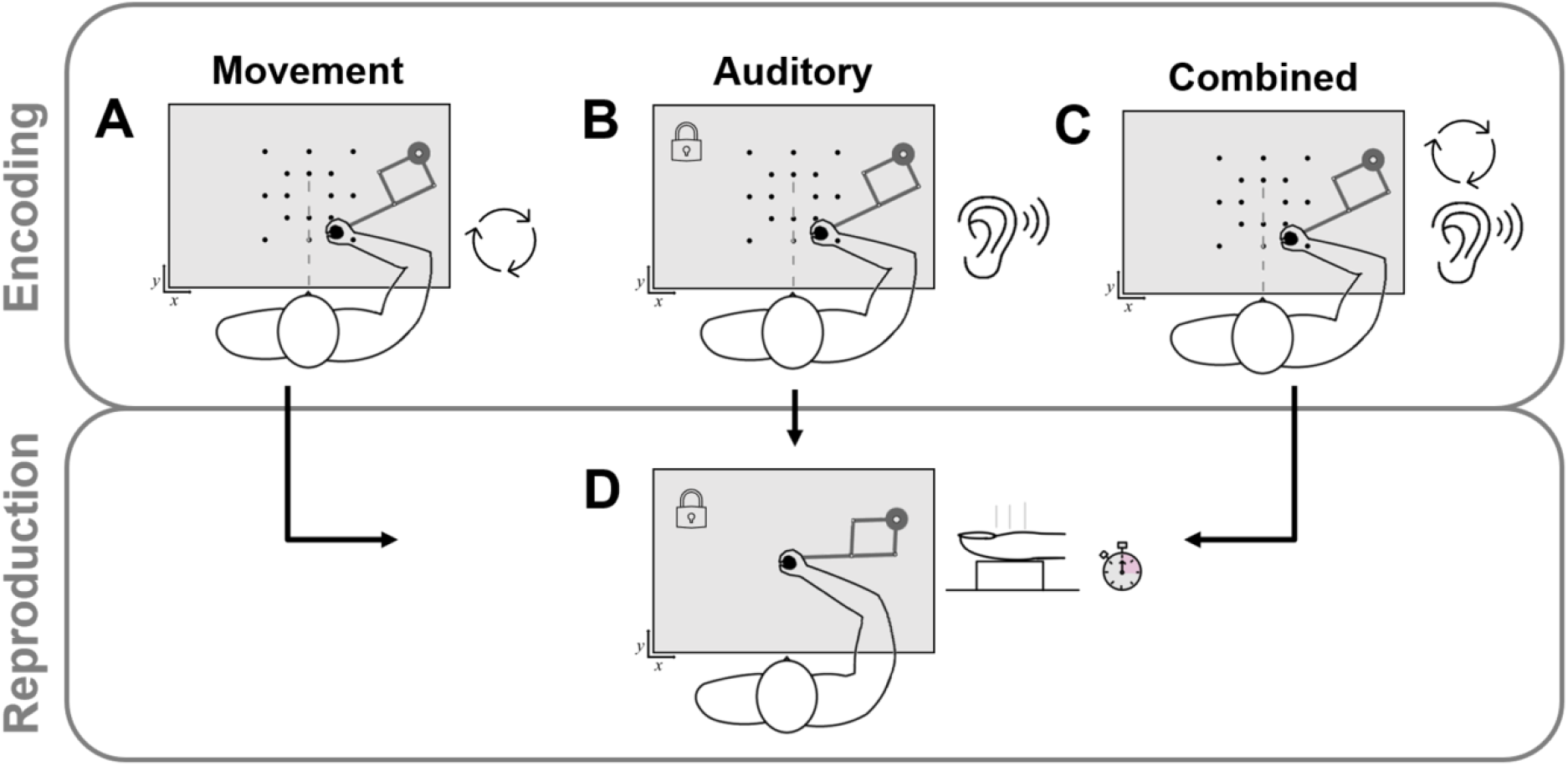
Task Schematic. (A) Movement-only trial: The robotic arm guided subjects to a random location (dots not visible during task), and the participant moved freely until interrupted by a “brake”. (B) Auditory-only trial: The robot handle was locked in random location, and the participant listened to an auditory tone. (C) Combined movement-auditory trial: The robotic arm was guided to a random location, and the participant moved freely and listened to the auditory tone until the “brake” was applied along with the tone offset. (D) Reproduction phase: the interval was estimated by holding and releasing a button attached to the handle.

### Analysis

Robotic arm manipulandum data was sampled at 1000 Hz to produce vectors for position, velocity, force, and other movement parameters over the course of time for each trial. Trials were excluded if reproduced times fell outside three deviations from the mean (<1% of trials excluded). Additionally, trials with movement (movement-only and combined) were excluded if the stop latency after the brake was applied fell outside three scaled absolute deviations from the median (2.9% of trials excluded).

Our first goal was to investigate the relationship between duration and reproduced time to determine timing accuracy. Importantly, movement-only and combined trials were analyzed with respect to time spent moving rather than the pre-specified duration. This is because in general, participants exhibited a short delay to respond to the movement brake, usually adding up to a few hundred milliseconds on most trials. We directly assessed constant error as a measure of accuracy, defined as the difference between the reproduced duration using the button press and the actual target duration. To measure performance, we applied a linear mixed model (LMM) design, in which response error was the predicted variable, encoded duration and condition were fixed effects, and subject was treated as a random effect.

Additionally, we examined the relationships between movement parameters and timing performance. Previous work has shown that arm movements covering a greater distance lead to longer perceived durations (De Kock et al., 2021; Wiener et al., 2019). Here, we extracted the movement distance for each trial (defined as the Euclidean distance traveled between duration onset and offset) and performed a Spearman partial correlation test between movement distance and reproduced time, controlling for target duration. Partial correlation values were calculated for each subject separately for movement and combined trial types; each was assessed with a one-sample t-test against a baseline of zero.

Statistical analyses were performed using R, JASP (http://www.jasp-stats.org), and Matlab. For accuracy and CV analyses, we report results from linear mixed models with subject as a random effect. Results are reported at a significance level of 0.05.

### Computational Modeling

To determine the sensitivity of each of the unisensory modality conditions, as well as the multisensory combined one, we employed a Bayesian Observer-Actor Model (Fig. 4) previously described by Remington and colleagues (Remington et al., 2018; Jazayeri and Shadlen, 2010) and used previously by our group (De Kock et al., 2021). In this model, sample durations (t_s_) are inferred as draws from noisy measurement distributions (t_m_) that scale in width according to the length of the presented interval. These measurements, when perceived, may be offset from veridical estimates as a result of perceptual bias or other outside forces (b). Due to the noise in the measurement process, the brain combines the perceived measurement with the prior distribution of presented intervals in a statistically optimal manner to produce a posterior estimate of time (t_e_. The mean of the posterior distribution is then, in turn, used to guide the reproduced interval (t_p_), corrupted by production noise (p). The resulting fits to this model thus produce an estimate of the measurement noise (m), the production noise (p), and the offset shift in perceived duration (b). Note that the offset term is also similar to that employed for other reproduction tasks as a shift parameter (Petzschner and Glasauer, 2011). Additionally, the prior used for the model can be either uniform or Gaussian in shape (Cicchini et al., 2012), with implications for how these are combined; uniform priors are characterized by the range of intervals presented, whereas gaussian priors are centered on the average duration presented, with a width dependent on their precision. We chose here to model the prior as a Gaussian, as each individual subject will have experienced a slightly different set of intervals during the estimation phase for each of the three conditions. This is because in the two movement conditions the offsets were variable from trial-to-trial. As such, we modeled the width of individual subject priors to match the width of encoded intervals for each subject (*σ*t_s_). Model fits were conducted by minimizing the negative log-likelihood of subject responses given the sample values using Matlab’s *fminsearch* function.

To determine if subjects combined auditory and movement modalities in an optimal manner, consistent with cue combination, we used outputs of the Bayesian model to compare between unisensory and multisensory conditions. Specifically, cue combination predictions that the multisensory combination *σ*_C_ of two unisensory estimates (auditory *σ*_A_ and movement *σ*_M_) when modeled as Gaussians, should equal:

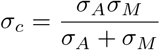

That is, the combined width is the product of the unisensory widths divided by their sum. Since the measurement widths were modeled as Gaussians in our Bayesian model, the unisensory widths from model fits can be used in the above equation to produce an estimate of the predicted width. Further, as the model also provides a width for the multisensory, combined condition, we can compare the width observed with that predicted by cue combination (Alais & Burr, 2019; Hartcher-O’Brien et al., 2014). If the predicted and observed widths match, then subjects combined unisensory estimates optimally, whereas larger observed widths than predicted indicate sub-optimal combination (Rahnev & Denison, 2018).

## Results

The temporal reproduction group data are illustrated in Fig. 2. As described above, movement-only and combined trials were analyzed with respect to time spent moving, which we defined as the predetermined interval (1000, 1500, 2000, 2500, 3000, 3500, 4000 ms) plus the “stop latency” (Fig. 2D), defined as the time it took participants to stop movement after the brake was applied. We used a linear mixed effects model with trial type (movement and combined levels only) and duration as fixed effects and subject as a random effect to test for stop latency differences related to trial type or duration. There were no significant main effects of trial type [F(1,2650.03)=.056, p=.812] or duration [F(1,2650.36)=.014, p=.906], and no significant interaction [F(1,2650.01)=1.622, p=.203]. In the rest of the figure panels, the target durations for movement and combined trials include these stop latencies, which are shown with horizontal error bars to denote the standard error (Fig. 2A-C).

**Figure 2:**
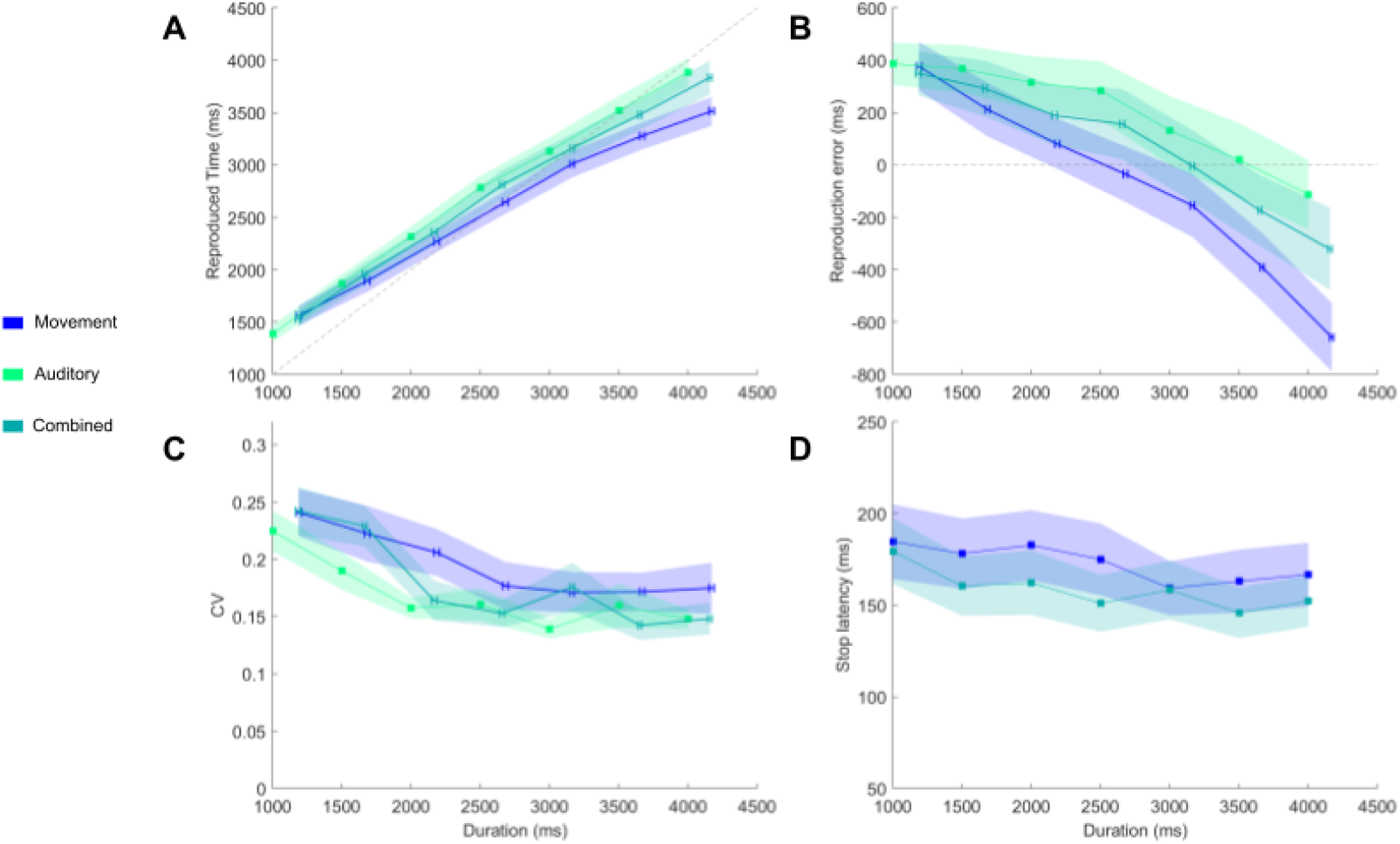
Temporal reproduction results. (A) Reproduction performance plotted as a function of target duration and trial type. Shaded vertical error bars represent reproduction standard error, and horizontal bars represent standard error of time spent moving for movement and combined trials. (B) Constant error values (reproduced duration - target duration) as a function of target duration and trial type. (C) Coefficient of variation (std/mean) of reproduction time as a function of target duration and trial type. (D) Stop latencies of movement and combined trials when the movement brake was applied. These values plus the fixed target durations (X axis) determined target durations for trials with movement.

We next employed a linear mixed effects model with trial type and duration as fixed effects and subject as a random effect to characterize how these variables affected reproduced duration. We did this analysis using constant error values (reproduced duration - target duration) rather than raw reproduction values as the dependent variable, as they represent the same underlying data but provide information about the direction and magnitude of errors (Fig. 2b). Model terms were nested using the Satterthwaite method. The model revealed a significant interaction of duration and trial type [F(2,4045.04)=30.577, p<.001], and main effects of trial type [F(2,4045.05)=5.512, p=.004] and duration [F(1,4045.15)=821.134, p<.001]. We examined the estimated marginal means and contrasts of trial types with Holm-adjusted p-values, which revealed a significant difference between auditory (EMM = 180.185, CI[-25.003, 385.374]) and movement (EMM = −51.534, CI[-256.860, 153.791]) trials (Contrast = 231.720, SE = 20.915, p < .001), combined (EMM = 81.551, CI[-123.649, 286.751]) and auditory trials (Contrast = −98.635, SE = 20.592, p < .001), and finally, between combined and movement trials (Contrast = 133.085, SE = 20.940, p < .001). We also compared the estimated marginal means to zero and did not find a significant result for any of the trial types (p_auditory_ = .085, p_movement_ = .623, p_combined_ = .436).

Our next analysis focused on the error slopes produced by the different trial types. This analysis reveals the degree of central tendency (i.e., attraction of estimates towards the mean). A significant slope difference was found between all trial types pairs (Auditory - Movement = .161, SE = .021, p < .001; Combined - Auditory = −.061, SE = .021, p = .003; Combined - Movement = .101, SE = .021, p < .001). Movement trials exhibited the lowest slope and auditory trials exhibited the highest slope.

Next, we examined the coefficient of variation (CV) as a measure of reproduction precision across durations and trial types. We employed a linear mixed model with trial type and duration as fixed effects and subject as a random effect, and found that the CV varied as a function of duration [F(1, 395.18)=64.227, p < .001] but not trial type [F(2, 395.05)=2.575, p = .077]. The interaction of trial type and duration was not significant [F(2, 395.05)=.893, p = .410].

We were additionally interested in the relationship between movement parameters and reproduced time, particularly for the combined trial type. We assessed this effect by performing subject-level partial correlations between Euclidean movement distance during duration encoding and the subsequent reproduced duration, controlling for target duration. The distribution of individual correlation coefficients is displayed in Fig. 3. Using a one-sample t-test, we found that the values were distributed significantly above zero for both motor [t(19)=2.974, p=0.008, Cohen’s D = 0.665] and combined [t(19)=4.278, p<0.001, D=0.957] conditions, indicating a positive relationship between movement distance and reproduced duration, and replicating prior work that movement distances are associated with longer estimated durations (Wiener et al. 2019; De Kock et al. 2021a).

**Figure 3:**
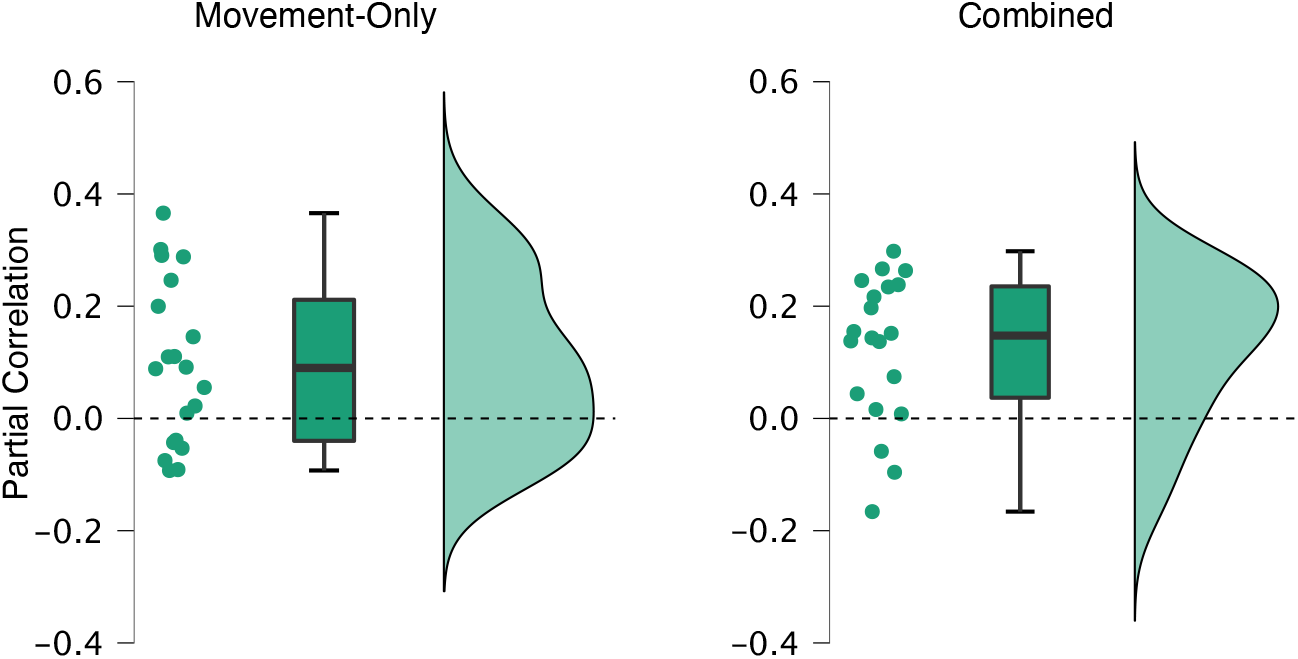
Effects of movement on duration estimates. Spearman partial correlation coefficients are displayed for each subject as raincloud plots in both the unisensory, movement-only condition and the multisensory, combined condition. For each subject, partial correlations were calculated as the association between single-trial reproduced durations and the movement length during the encoding phase, while controlling for duration. On average, the correlation coefficients in both groups were greater significantly greater than zero.

### Cue-Combination

To examine the impact of modality on time estimates, we further fit reproduced durations with a Bayesian Observer-Actor model. The results of our model fits provided estimates of the measurement noise (*m)*, production noise (*p*), and offset (*b*) for each modality. Repeated-measures ANOVAs for each parameter revealed only a main effect of condition for measurement noise [*F* (2,38)=2.133, *p*=0.039], with all other parameters non-significant (all *p>*0.05). For the noise parameter, measurement noise scores were significantly lower for the combined condition compared to movement [*t* (19)=−2.118, *p*=0.048], but not auditory intervals [*t* (19)=−1.682, *p*=0.109] (Fig. 4). Due to our a-priori hypothesis that the combined multisensory estimates would be better than both unisensory estimates, we averaged auditory and movement measurement widths and compared them to the combined measurement noise, where a significant difference was observed [*t* (19)=− 2.15, *p*=0.045].

**Figure 4:**
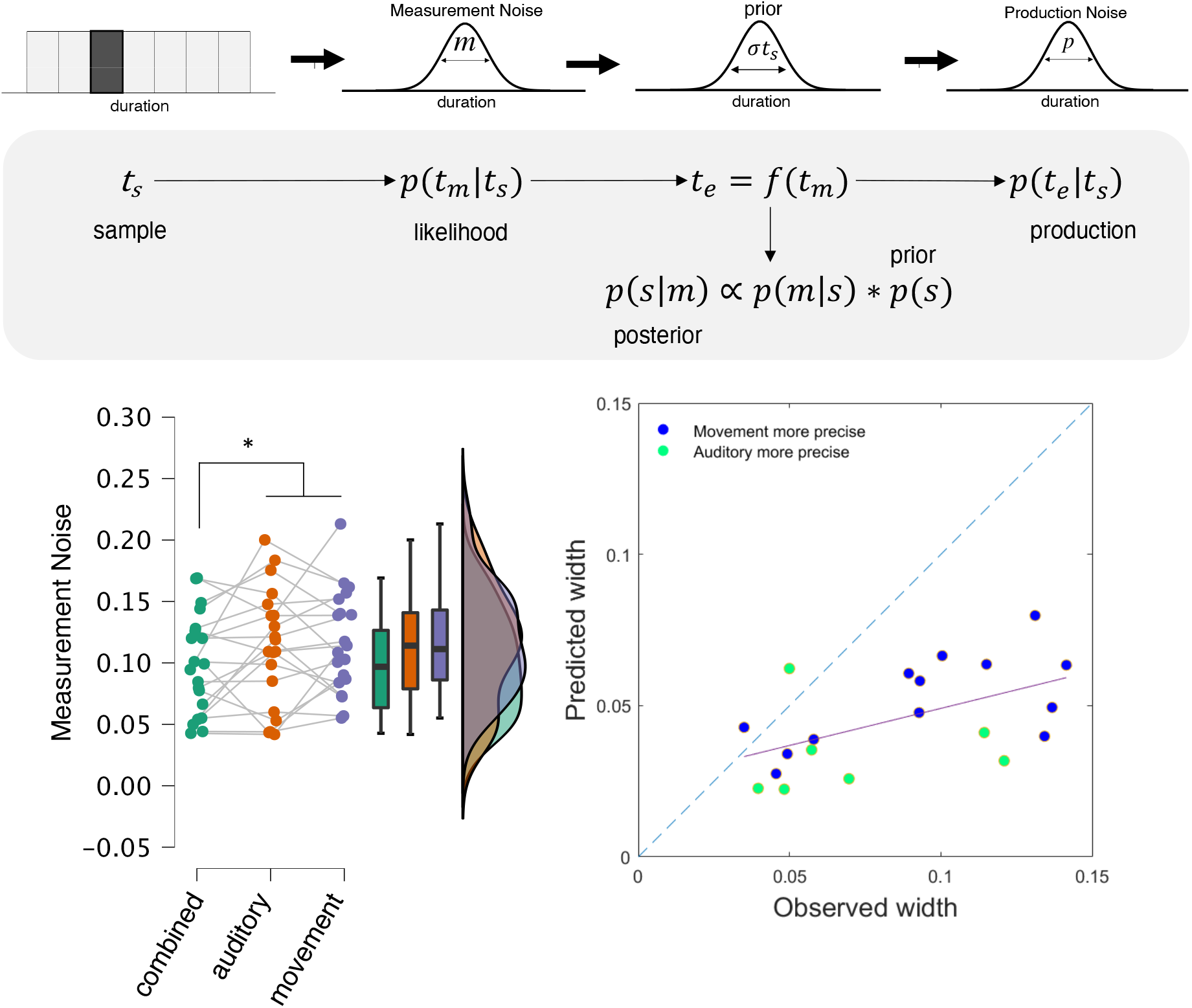
Bayesian Observer-Actor Model for time estimates. Top: Schematic of the model, in which a sample interval presented on a given trial *t_s_* is perceived with some measurement noise from a Gaussian distribution with width *m*. This estimate is then combined with a prior distribution of previously-experienced intervals, also characterized as a Gaussian, with width *σt_s_* to form a posterior estimate *t_e_*. During reproduction, the posterior estimate is further corrupted by motor production noise as a Gaussian distribution with width *p*. For model fits, *m* and *p* were set as free parameters, whereas *σt_s_* was measured directly from the experienced sample intervals for each subject. Bottom Left: model fits for the measurement noise width (*m*) for all three modality conditions. We observed that the multisensory, combined measurement noise was significantly lower than both the unisensory auditory and movement conditions, together. Bottom Right: Scatterplot comparing the observed multisensory width to the predicted width given by combining both of the unisensory widths together via cue combination. Here, we observed that the observed widths fell on average below the predicted width, yet were also significantly correlated. Individual points are colored based on which of the unisensory widths was smaller (more precise); no differences were found between either of these groups.

To compare with the predictions of the cue combination model, we used the unisensory measurement noise widths to generate a predicted width for their optimal combination. That is, the product of the unisensory widths dived by their sum. Here, we observed that the predicted widths were on average lower than the observed widths from the combined multisensory condition [*t* (19)=6.111, *p*<0.001] (Fig. 4). Additionally, we observed a strong correlation between observed and predicted widths [*r* (18)=0.654, *p*=0.002], indicating that the cue combination equation provided a good prediction of multisensory noise, even if the observed estimates were sub-optimal. As a further observation, we noted that subjects with lower multisensory noise estimates were closer to the optimal prediction, a finding we confirmed quantitatively through the correlation of multisensory widths with the difference between those widths and the predicted estimate [*r* (18)=−0.883, *p*<0.001]. As a final check, we compared subjects who exhibited smaller unisensory widths for one modality (e.g., auditory) over the other (e.g. movement). No differences were observed between subjects with greater precision in either modality, for either the multisensory combined estimates [*t* (18)=0.765, *p*=0.454], or the difference from optimality [*t* (18)=−0.528, *p*=0.604] indicating that improved estimates for neither unisensory modality conveyed a special benefit in cue combination.

## Discussion

We administered a temporal reproduction experiment in which we tested timing performance for auditory and motor timing, and both simultaneously. Measuring these two sources of information allowed us to assess how self-movements are judged in comparison to auditory tones, and shed light on the computational mechanisms that drive movement-related improvements previously observed during perceptual timing tasks ((Wiener et al., 2019); (De Kock et al., 2021)b). We found that modality significantly impacted reproduction performance, such that motor trials resulted in shorter estimates than auditory trials. Critically, combined trials were estimated most accurately, suggesting that the natural biases introduced by auditory and motor intervals oppose each other, but “work together” to form the most veridical estimate based on the information available. Additionally, our results fit into a framework of Bayesian cue combination in which multimodal interval measurements are more precise than unisensory measurements ((De Kock et al., 2021)b). We also found that the reproduction slope was lowest for the movement condition, indicating a greater degree of central tendency compared to the other conditions. An unanswered question is the extent to which this reflects intrinsic properties of movement interval timing, such as a greater reliance on an internalized prior distribution (Cicchini et al., 2012) or a general susceptibility towards underestimation as target durations increase.

These results corroborate precious accounts of movement enhancing cross-modal timing ((Wiener et al., 2019); (Iordanescu et al., 2013); (Carlini & French, 2014); (Manning & Schutz, 2013); (Safaie et al., 2020)). However, it is of interest whether this is accomplished in an optimal fashion. According to the Bayesian cue combination framework, we predicted that the measurement noise (i.e., distribution width) of the combined condition would be lower than for unimodal conditions. The model results indicated that the measurement noise of the combined condition was significantly lower than for movement and auditory conditions together. While we did not observe meaningful differences in coefficient of variation (an index of variability that is often tied to precision; (Brown, 1997); (Lewis & Miall, 2009)) between trial types, we suggest that the measurement noise parameter is a more useful indicator of precision in our experiment given that the target durations were not fixed for trials that involved movement, and further that this measure attempts to remove motor production noise. We next compared the observed measurement noise to the model prediction that would indicate optimally combined estimates. The observed and predicted values were significantly positively correlated, although we note that participants generally combined sub-optimally. This pattern was not dependent on which unimodal condition was more precise for individual participants. The model also predicts that during optimal multimodal timing, the mean reproduction estimate should gravitate towards the more precise modality; however, given that we did not find overall differences in unimodal timing precision across participants and performance was generally sub-optimal, this prediction was challenging to test in the current paradigm. Remarkably, this lack of a difference highlights that movement timing is at least as precise as auditory timing, which until now has been documented as the most precisely timed modality ((Wiener et al., 2014; Jones et al., 2009); (Burr et al., 2009)). Future work may assess this prediction more closely where larger differences between unimodal conditions exist (e.g., visual timing).

We have described some computational principles by which motor and sensory information may be combined for a more precise multimodal estimate. These perspectives are strengthened by discussing their relation to neural mechanisms. There is a great degree of functional overlap between motor and timing activity in brain regions considered vital to motor control, with greater representation of supra-second intervals in cortical regions and sub-second intervals in subcortical regions like the cerebellum ((Nani et al., 2019); (Wiener et al., 2010)). The supplementary motor area (SMA) stands out as a region of interest, as it is activated across a wide range of timing tasks and encodes time intervals in neurons organized along a rostrocaudal “chronotopic” gradient (Protopapa et al., 2019). Additionally, the SMA exists within a larger cortico-thalamic-basal ganglia timing circuit (Merchant & Yarrow, 2016) that encodes intervals, integrates multiple sensory inputs (Nagy et al., 2006), and sends predictive signals to sensory areas.

Using these insights, we have outlined two possibilities to describe the neural implementation of movement-related timing effects ((De Kock et al., 2021)b). The first possibility, Feedforward Enhancement, posits that these effects are instantiated in motor circuits and sharpen duration measurements via corollary feedback to motor regions like the SMA, thereby sharpening the tuning of duration-selective neurons. The SMA is equipped to respond to these signals via white matter connections with the primary motor cortex, basal ganglia and spinal cord (Vergani et al., 2014), all of which require a high degree of temporal coordination during movement control. Additionally, deep reinforcement learning agents with feedforward modules (and no recurrence) successfully learn to produce temporal intervals by generating stereotyped trajectories in their environments (Deverett et al., 2019).

The second possibility, Active Sensing (Schroeder et al., 2010), proposes that motor activity acts on earlier sensory regions to enhance cross-modal temporal measurements. Outside of the timing domain, this is a well-established process. For example, auditory perception is enhanced when a sound is triggered by an action (Myers et al., 2020), and accordingly, motor preparation has been found to elicit responses in the auditory cortex (Gale et al., 2021). There are also several examples tying motor activity to changes in visual processing ((Yon et al., 2018); (Tomassini et al., 2020); (Saleem et al., 2013); (Niell & Stryker, 2010); (Cao & Händel, 2019)). The Active Sensing hypothesis is compatible with prior research on multisensory integration, as it essentially describes the convergence of multiple signals to shape neural computations. A classic example is the superior colliculus which integrates visual, auditory, and sensorimotor signals (among others) to guide eye and head movements ((Stein, 1998); (Distler & Hoffmann, 2015)). These signals are not localized to dedicated hubs, but rather occur throughout the neocortex (Ghazanfar & Schroeder, 2006), and have been proposed to reflect generalizable “canonical operations” (e.g., divisive normalization and oscillatory phase resets) when integrating a diverse range of inputs – including from motor circuits (Van Atteveldt et al., 2014). For example, saccade onsets elicit time-locked local field potential changes in primary visual cortex (Ito et al., 2011). The Active Sensing hypothesis has empirical support from several lines of research, and can build upon known neural integration mechanisms – more generalizable than previously thought – to shed light on how movement can improve timing.

Beyond basic neuroscience, this work has implications in clinical disorders associated with motor and timing deficits such as Parkinson’s Disease and Huntington’s Disease ((Avanzino et al., 2016); (Singh et al., 2021); (Merchant et al., 2008); (Cope et al., 2014)). These parallel deficits are not restricted to movement disorders, but occur in psychiatric or neurodevelopmental conditions such as schizophrenia and attention deficit hyperactivity disorder ((Walther & Strik, 2012); (Yang et al., 2007)). Motor training has shown some usefulness in rehabilitation and symptom management; Parkinson’s patients have been found to reduce their gait variability when exposed to rhythmic auditory stimuli that can adaptively synchronize with their steps (Miyake, 2009). In stroke patients, fine motor skills are re-learned more effectively with musical motor training than functional motor training (Schneider et al., 2010). Thus, many benefits of movement training rely strongly on integrating relevant sensory information, and based on this evidence, interactions between motor and auditory interval timing may be a promising avenue to explore in the treatment and diagnosis of movement disorders.

In conclusion, while converging evidence suggests a powerful role of movement in shaping time perception ((De Kock et al., 2021)b; (Balasubramaniam et al., 2021)), studies have focused primarily on perceptual timing with movement as an added component without isolating how movements are consciously timed on their own. Our experiment provided the distinct advantage of isolating and comparing timing in movement and auditory modalities, in addition to testing predictions about their integration under a Bayesian cue combination framework. We found that multisensory timing was superior to unisensory timing as reflected in the higher accuracy of combined estimates, and additionally, measurement noise of combined estimates reflected at least some degree of optimal cue combination (with the caveat that participants often perform sub-optimally compared to computational models). This study thus addresses a prior gap in knowledge where consciously-timed self-movement were not well understood, especially in relation to timing in other modalities. We also expanded on a growing body of research on movement-timing effects, first by describing potential computational mechanisms that drive them, and ways they may be instantiated in neural circuits.

